# A Systematic Analysis of Atomic Protein-Ligand Interactions in the PDB

**DOI:** 10.1101/136440

**Authors:** Renato Ferreira de Freitas, Matthieu Schapira

## Abstract

As the protein databank (PDB) recently passed the cap of 123,456 structures, it stands more than ever as an important resource not only to analyze structural features of specific biological systems, but also to study the prevalence of structural patterns observed in a large body of unrelated structures, that may reflect rules governing protein folding or molecular recognition. Here, we compiled a list of 11,016 unique structures of small-molecule ligands bound to proteins – 6,444 of which have experimental binding affinity - representing 750,873 protein-ligand atomic interactions, and analyzed the frequency, geometry and impact of each interaction type. We find that hydrophobic interactions are generally enriched in high-efficiency ligands, but polar interactions are over-represented in fragment inhibitors. While most observations extracted from the PDB will be familiar to seasoned medicinal chemists, less expected findings, such as the high number of C–H **…**O hydrogen bonds or the relatively frequent amide-π stacking between the backbone amide of proteins and aromatic rings of ligands, uncover underused ligand design strategies.

## Introduction

Significant progress in high-throughput X-ray crystallography^1,2^ combined with advances in structural genomics^3–5^ have led to an explosion in the number of structures publicly available in the Protein Data Bank (PDB).^6^ At the time this manuscript was written, more than 123,456 structures had been deposited in the PDB^6^, including 76,056 protein-small molecule complexes, of which 13,000 have a reported binding potency.^7,8^ This large body of data contains important information on the nature, geometry, and frequency of atomic interactions that drive potent binding between small molecule ligands and their receptors. Systematic analysis of this data will lead to a better appreciation of intermolecular interactions between proteins and their ligands and can inform structure-based design and optimization of drugs.^9^

Several approaches have been developed for large-scale analysis of protein–small molecule interactions, such as SuperStar, or that implemented to build the Relibase database.^10,11^ PDBeMotif^12^ and the recently published PELIKAN^13^ are two examples of free tools that can search for patterns in large collections of protein-ligand interfaces. Structural interaction fingerprints (SIF)^9^ are another method of representing and analyzing 3D protein-ligand interactions where the presence or absence of interactions between distinct residues and ligand atoms are represented as bit strings that can be compared rapidly.^14^ In addition, there has been an increase in the number of free tools to fully automate the detection and visualization of relevant non-covalent protein–ligand contacts in 3D structures.^15–17^

A statistical analysis of the nature, geometry and frequency of atomic interactions between small molecule ligands and their receptors in the PDB could inform the rational optimization of chemical series, help in the interpretation of difficult SAR, and serve as a knowledge-base for the improvement of scoring functions used in virtual screening. To the best of our knowledge, such public resource is currently missing.

Here, we analyze the frequency of common atomic interactions between protein and small molecules observed in the PDB. We find that some interactions occur more frequently in fragments than drug-like compounds, or in high-efficiency ligands than low-efficiency ligands. We next review in detail each of the most frequent interactions and use matched molecular pairs to illustrate the impact of these atomic interactions on binding affinity.

## Most Frequent Protein-Ligand Atomic Interactions

We extracted from the PDB all X-ray structures of small-molecules in complex with proteins, with a resolution ≤ 2.5 Å, resulting in a collection of 11,016 complexes (only one structure was kept for each ligand). This collection contained 750,873 ligand-protein atom pairs, where a pair of atoms is defined as two atoms separated by 4Å or less. The top-100 most frequent ligand-protein atom pairs (Supplementary Table 1) can be clustered into seven interaction types (Figure 1). Among the most frequently observed are interactions that are well known and widely used in ligand design such as hydrophobic contacts, hydrogen bonds and π–stacking.^18,19^ These are followed by weak hydrogen bonds, salt bridges, amide stacking, and cation-π interactions.

**Figure 1.**
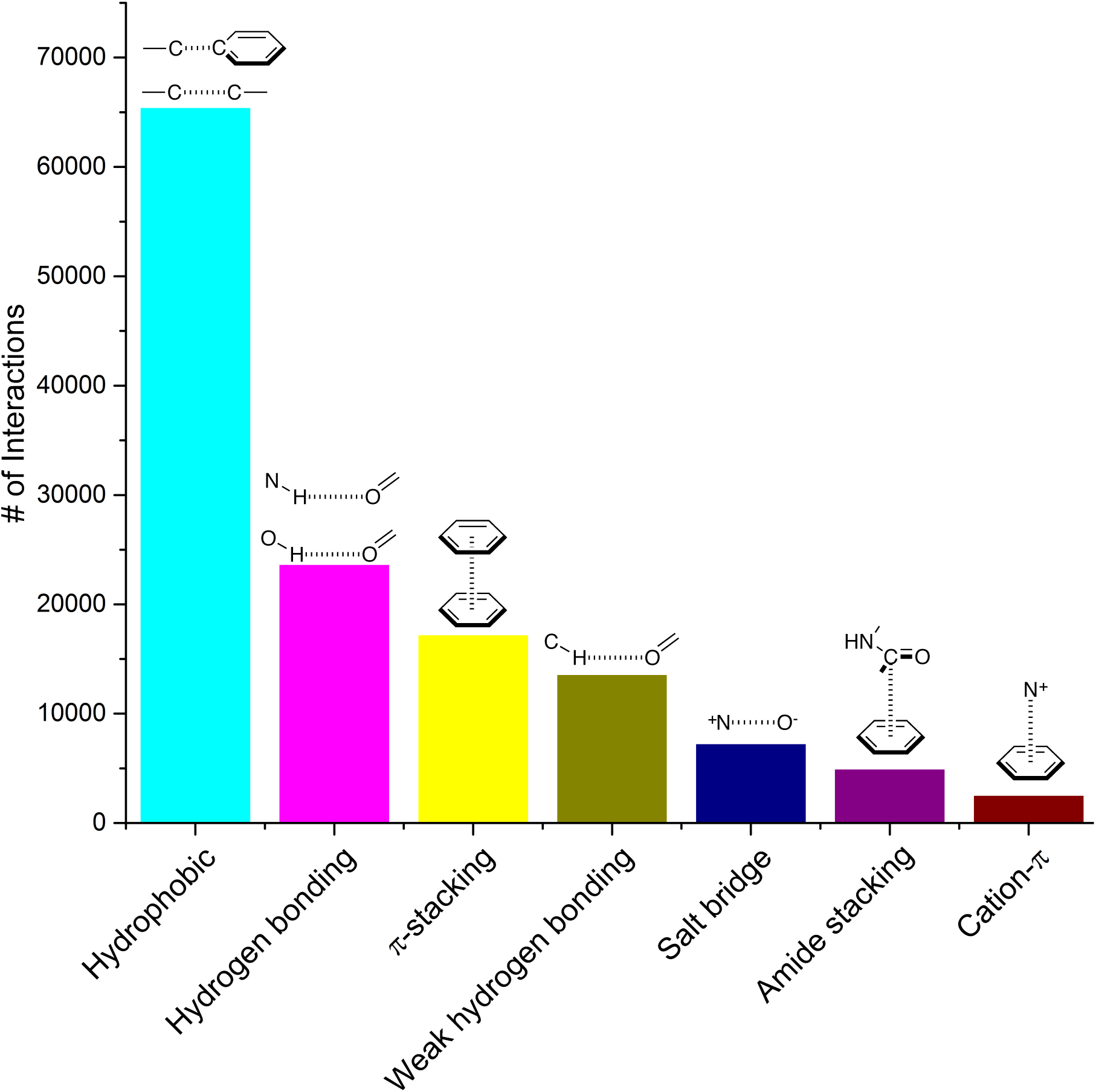
Frequency distribution of the most common non-covalent interactions observed in protein-ligands extracted from the PDB.

We next asked whether some interactions types were more frequently observed in high-efficiency ligands. Experimental binding affinity for 6444 protein-ligands in the PDB were retrieved from the PDBbind database^7,8^, and a fit quality (FQ) score - a size-adjusted calculation of ligand efficiency - was used to evaluate how optimally a ligand binds relative to other ligands of any size.^20^ The frequency of each interaction type was calculated in the 1500 protein-ligand complexes with the best FQ score (FQ > 0.81) and the 1500 complexes with the worst FQ scores (FQ < 0.54) (Figure 2a).

**Figure 2.**
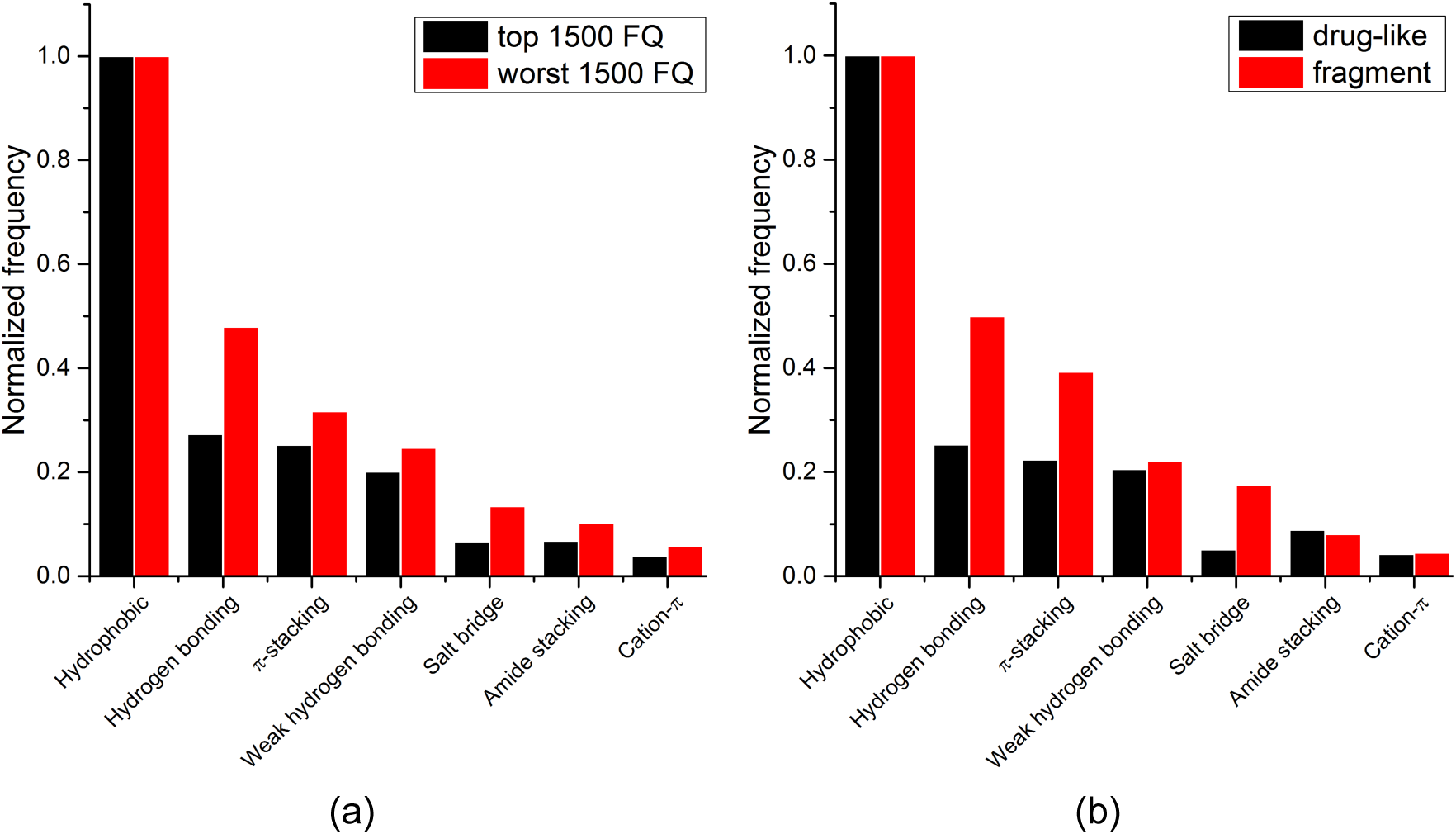
Relative frequency distribution of the most common non-covalent interactions observed in: (a) ligands with high vs. low ligand efficiency; (b) fragments vs. drug-like compounds.

We find that hydrophobic interactions are more frequent in high-efficiency ligands. In particular, the relative contribution of hydrogen bonds is reduced from 48% to 27% of that of hydrophobic contacts in efficient binders, and the contribution of salt bridges is more than halved, from 13% to 7%. This observation probably reflects the fact that most ligands in the PDB are the product of lead optimization strategies that aim at increasing the number of favorable hydrophobic interactions, which is less challenging than optimizing directionality-constrained hydrogen-bonds (discussed in more details in a later section).^21^ We also find that efficient ligands are more hydrophobic, as the median number of heavy atoms and logD (ChemAxon) for compounds with high FQ are 27 and 1.7 respectively, and 21 and 0.2 respectively for compounds with low FQ. Both groups showed similar profiles for other properties like polar surface area (median PSA: 95.3 *vs*. Å^2^), hydrogen bond acceptors (median HBA: 5 for both), and hydrogen bond donors (median HBD: 2 for both). Taken together these results show that small-molecule ligands that bind their target with high efficiency are more hydrophobic, and that hydrophobic interactions are a driving factor for increased ligand efficiency.

Since fragments are typically binding their targets with higher efficiency than larger ligands, we asked whether hydrophobic interactions were also more frequent in protein-fragment complexes. The frequency of each interaction type was calculated for two random groups of 1500 hundred protein-ligand complexes, one with fragment molecules (HA ≤ 20), the other with drug-like molecules (30 ≤ HA ≤ 50) (Figure 2b). Unlike high-efficiency ligands, we find that protein-fragment complexes are enriched in polar interactions: the frequency of hydrogen bond is doubled, from 25% to 50%, compared to that of hydrophobic contacts, and the frequency of electrostatic interactions is multiplied by three, from 5% to 17%. To compensate for their low number, interactions made by fragments need to be highly efficient.^22^ We note that electrostatic interactions define maximum efficiency of ligand binding.^23^ Similarly, though to a lesser extent, optimal hydrogen-bonds are energetically more favorable than hydrophobic contacts.^24^ The observed enrichment in polar interactions reflects the fact that fragments are freer than chemical moieties of larger compounds to adopt binding poses that will optimally satisfy the geometric constraints of high-efficiency interactions, such as electrostatic or hydrogen-bonds.^25^

Together, these results show that fragments are using polar interactions to gain maximum binding efficiency from a limited number of interactions, but as small-molecule ligands are optimized, geometric constraints associated with polar bonds are more challenging to satisfy, and the contribution of hydrophobic interactions increases. To gain further insight, we next analyzed in detail the composition, geometry, frequency, protein side-chain preference, and impact towards binding affinity of each protein-ligand interaction type in the PDB.

## Specific Intermolecular Interactions

### Hydrophobic interactions

From our analysis, hydrophobic contacts are by far the most common interactions in drug-receptor complexes, totalizing 66772 contacts between a carbon and a carbon, halogen or sulfur atom (the distance cut-off of 4.0 Å allows the implicit inclusion of hydrogen atoms) (Figure 1). Hydrophobic interactions were separated into five groups (Table S2). The most populated group is the one formed by an aliphatic carbon in the receptor and an aromatic carbon in the ligand, which alone accounts for more than 42000 interactions (Table S2). This is an indication that aromatic rings are prevalent in small molecule inhibitors. In fact, 76% of the marketed drugs contain one or more aromatic ring, with benzene being by far the most frequently encountered ring system.^26,27^ Not surprisingly, leucine, followed by valine, isoleucine and alanine side-chains are the most frequently engaged in hydrophobic interactions (Figure S2).

Contacts involving an aromatic or aliphatic carbon in the receptor and an aliphatic carbon in the ligand were observed in 8899 and 8974 instances, respectively (Table S2). We observed that aliphatic carbons were distributed mostly above or below the plane of the aromatic ring, rather than at the edge (Figure S3). Although we didn’t investigate further the directionality of these interactions we believed that some of them could be further classified as C-H**…**π interactions.^28^ These interactions occur ubiquitously in almost all proteins, making important contributions to biomolecular structures and ligand binding.^29,30^ Interactions involving an aliphatic or aromatic carbon in the protein and a chlorine or fluorine in the ligand were observed 5147 times (Table S2). While these atomic contacts are hydrophobic, a C–X**…**π component can also contribute to this type of interaction.^31,32^ Interactions involving a sulfur atom from the side chain of methionine and an aromatic carbon from the ligand were observed in 1309 complexes. Although methionine is classified as a hydrophobic residue, a recent study shows that the Met S**…**C(aro) interaction yields an additional stabilization energy of 1–1.5 kcal/mol compared with a purely hydrophobic interaction.^33^

Hydrophobic interactions are the main driving force in drug-receptor interactions. The benefit of burying a solvent-exposed methyl group on a ligand into a hydrophobic pocket of a protein is about 0.7 kcal/mol or a 3.2-fold increase in binding constant per methyl group.^34^ However the effect of replacing a hydrogen atom with a methyl group is highly context dependent, and potency losses are as common as gains. Ten-fold and 100-fold gains in potency are observed in 8% and 1% of cases respectively.^35,36^ For instance, addition of a single methyl group improves by 50 fold the potency of a tankyrase-2 (TNKS2) inhibitor.^37^ The added methyl group occupies a small hydrophobic cavity and potentially releases unfavorably bound water molecules (Figure 3b). In rare cases, the increase in potency due to the introduction of a “magic methyl” exceeds two orders of magnitude.^38^ This is generally due to the combined entropic effect of lowering the conformational penalty paid by the ligand upon binding, and the desolvation effect of burying the methyl group in a hydrophobic pocket.^35^

**Figure 3.**
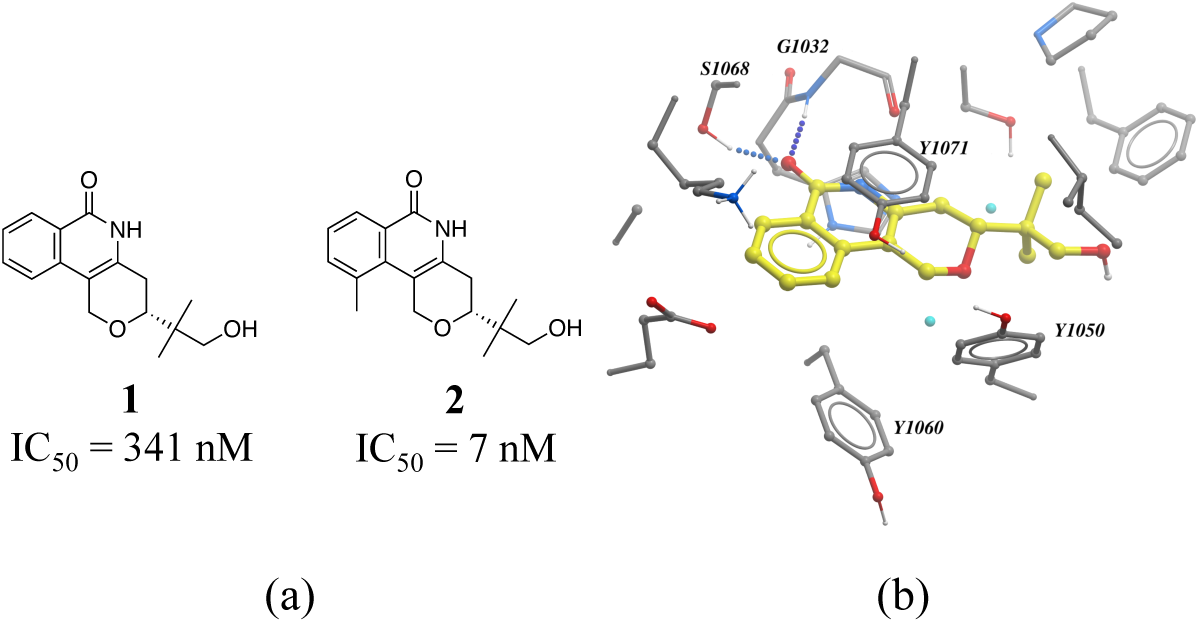
Magic methyl effect: (a) Chemical structure of two TNKS2 inhibitors; (b) crystal structure of **1** (carbon atoms in yellow) bound to TNKS2 (PDB: 5C5P). Oxygen atoms are in red (cyan for water molecules).

### Hydrogen bonds

We find that hydrogen bonds were the second most frequent type of interactions observed in our collection of protein-ligand complexes, with a total number of 24058 instances (Figure 1). N-H**…**O interactions were more frequent (15105 interactions) than O-H**…**O (8251 interactions) and N-H**…**N (333 interactions) (Table S2). Among the N-H**…**O interactions, the number of neutral and charged hydrogen bonds were almost equal (7554 *vs*. 7551, respectively). Proteins were more often hydrogen-bond donors than acceptors (9217 *vs*. 5888, respectively). Surprisingly, glycine was the most frequent hydrogen-bond acceptor, and the second most frequent donor (Figure S4). Arginines were engaged in more hydrogen-bonds than lysines, probably reflecting the presence of 3 nitrogen atoms in the guanidinium group of arginine side-chains (Figure S4). Among O-H**…**O interactions, charged hydrogen bonds (typically between an alcohol and a carboxylic acid) were 3 times more frequent than neutral ones, and ligands more often behaved as donors than acceptors (Table S2). The most common acceptors were aspartic acids in charged hydrogen bonds, and asparagine, glycine and glutamine in neutral interactions (Figure S5). Serine was the most usual donor (Figure S5).

We found that heavy atoms in N-H**…**O, N-H**…**N, and O-H**…**O hydrogen bonds were all separated by similar median distances of approximately 3.0 Å (Figure 4). This value is slightly higher (∼ 0.1-0.2 Å) than previously reported for hydrogen bonds between amide C=O and OH/NH.^39^ In addition, the median distances of neutral and charged hydrogen bonds were almost identical (0.1 Å difference, data not shown). The D-H**…**A angle usually peaked at 130-180°, and the preferred angle for N-H**…**O hydrogen bonds was around 180° (data not shown)

**Figure 4.**
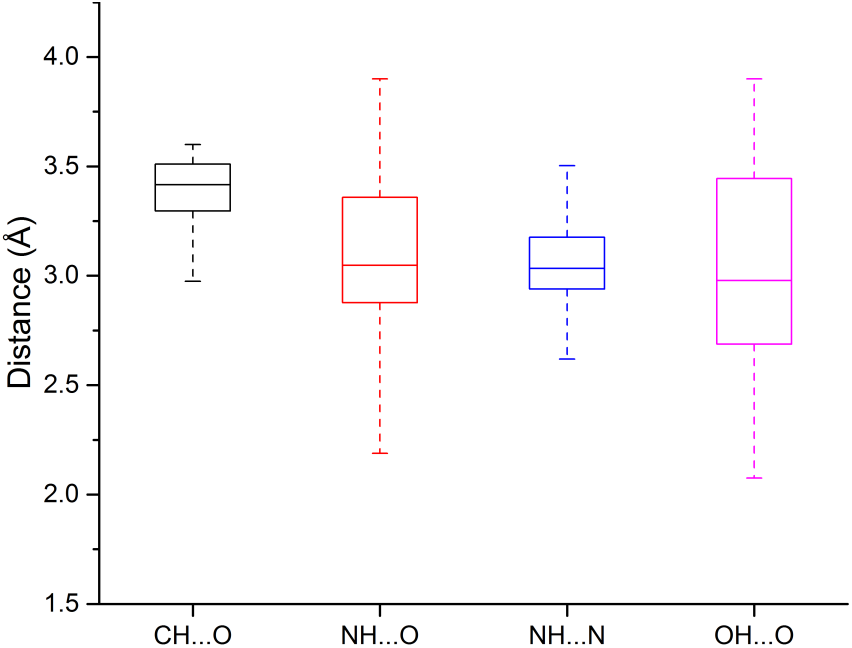
Box plots of hydrogen bond length distributions for the weak (C-H**…**O) and strong hydrogen bonds (N-H**…**O, N-H**…**N, O-H**…**O).

Hydrogen bonds are the prevailing directional intermolecular interactions in biological complexes^40^, and the predominant contribution to the specificity of molecular recognition.^41^ Hydrogen-bond donors and acceptors not only play important roles in DNA base-pairing^42^, protein folding^43,44^, and enzyme specificity^45^, but also in modulating the physicochemical properties - in particular the lipophilicity^46^ - of ligands, a key parameter in drug design. The free energy for hydrogen bonding can vary between -1.5 kcal/mol to -4.7 kcal/mol.^34^ However, the contribution of a hydrogen bond to binding can be very modest (or penalizing) if the new interaction formed does not outweigh the desolvation penalty upon ligand binding.^47^ Also, the contribution of a hydrogen bond is dependent on the local environment: a solvent-exposed hydrogen-bond contributes significantly less to net interaction energy than the same hydrogen-bond in a buried hydrophobic pocket.^48^ Consequently, optimizing hydrophobic interactions is generally considered easier than hydrogen bonds.^34^ In drug design, hydrogen bonds are exploited to gain specificity owing to their strict distance and geometric constraints.^49^

Among numerous examples, a series of potent thrombin inhibitors shows a remarkable increase in binding affinity (> 500-fold) through simple addition of hydrogen-donating ammonium group (Figure 5).^50^ In the crystal structure, the ammonium group forms a charge-assisted hydrogen bond with the carbonyl oxygen of Gly216 and surrounding waters.^51^

**Figure 5.**
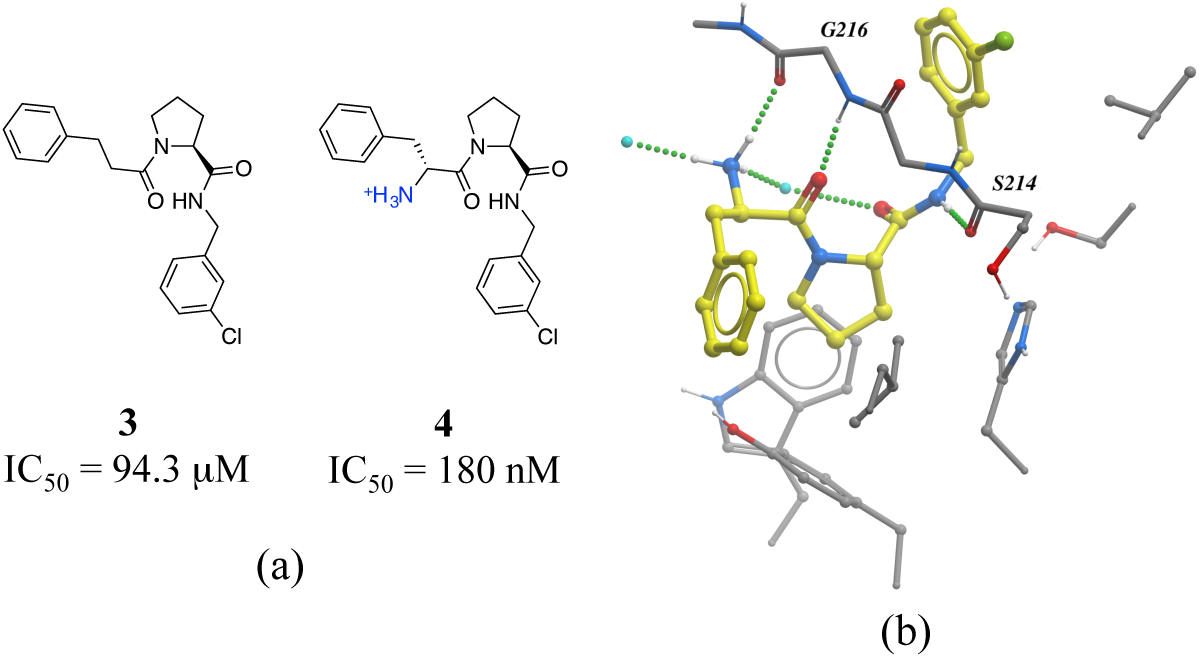
Effect of adding a hydrogen bond in a thrombin inhibitor: a) chemical structure of a pair of thrombin inhibitors; b) crystal structure of **4** (yellow carbons) in complex with thrombin (PDB: 2ZC9). Hydrogen bonds are displayed in dotted green lines and waters oxygens as cyan.

### π -stacking interactions

The third most frequent protein-ligand contacts in the PDB were aromatic interactions (Figure. 1). Interactions involving aromatic rings are ubiquitous in chemical and biological systems and can be considered a special case of hydrophobic interactions.^28^ We found that edge-to-face and face-to-face interactions were equiprobable (8704 and 8537 contacts respectively) (Table S2). This is in agreement with quantum mechanical calculations of the interaction energy of benzene dimers that predict the edge-to-face and parallel displaced face-to-face as being isoenergetic, and more stable than the eclipsed face-to-face π-stacking.^52^ Almost 50% of all π-stacking interactions are observed between the aromatic ring of phenylalanine and an aromatic ring in the ligand, followed by tyrosine (36.8%), tryptophan (8.7%) and histidine (5.1%) (Figure S6).

Interactions involving aromatic rings are major contributors to protein–ligand recognition and concomitantly to drug design.^28,53^ An example of the strong gain in binding affinity that can be obtained by forming a π-stacking interaction is illustrated in a series of soluble epoxide hydrolase (sEH) inhibitors.^54^ In the X-ray cocrystal structure of human sEH and **6** (IC_50_ = 7 nM), the phenyl ring is positioned to allow π-stacking interaction with H524 (Figure 6b), while an analog (**5**, IC_50_ = 700 nM) without the phenyl ring is l00-fold less potent. While π-stacking interaction can increase the binding affinity of the inhibitor for its target, it has been pointed out that reducing the number of aromatic rings of a molecule might improve its physicochemical properties, such as solubility.^55,56^

**Figure 6.**
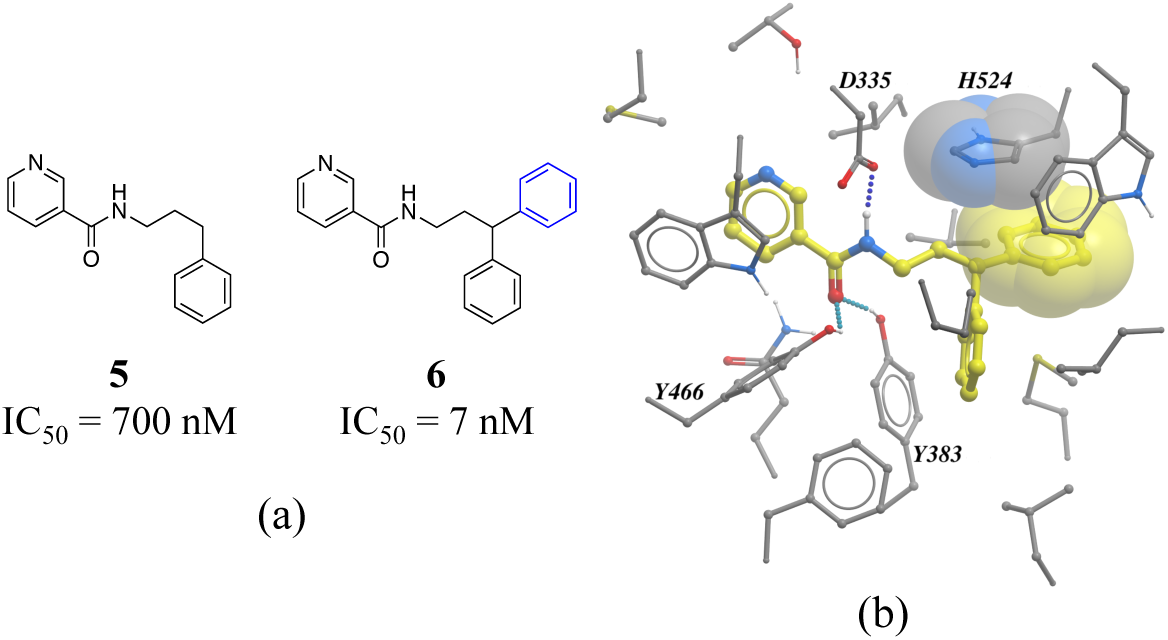
a) Chemical structure of two inhibitors of human sEH; b) X-ray cocrystal structure of human sEH and **6** (yellow carbons, PDB: 3I1Y), the phenyl ring is positioned to allow a π-stacking interaction with H524. Hydrogen bonds are displayed in dotted green lines.

### Weak hydrogen bonds

The fourth most frequent interactions (13600 contacts) were C-H**…**O hydrogen bonds, the existence of which is well documented (Figure 1).^57,58^ When the interacting carbon was aromatic, protein oxygens were found to be acceptors much more often than ligand oxygen atoms (4927 *vs.* 708 interactions, Table S2). This simply reflects the fact that most ligands have aromatic rings, while most side-chains don’t. Glycine, aspartic acid and glutamic acid were always the most frequent acceptors in C-H**…**O interactions, while leucine was the most frequent donor (Figure S7).

The median distance of the C-H**…**O hydrogen bonding was 3.4 Å, which is 0.4 Å longer than traditional hydrogen bonds (N-H**…**O, N-H**…**N, O-H**…**O), and distances separating the two heavy atoms were rarely lower than 3.2 Å (Figure 3). The angle distribution of C-H**…**O interactions peaked around 130° (data not shown), which is in agreement with previous work.^59^

The existence of weak hydrogen bonds has been extensively analyzed and reviewed.^60–63^ Calculations indicate that the magnitude of the C_α_-H**…**O=C interactions are about one-half the strength of an NH**…**O=C hydrogen bond.^64^ In addition, an analysis of protein-ligand complexes revealed that C_α_-H**…**O hydrogen should be better interpreted as secondary interactions, as they are frequently accompanied by bifurcated N-H**…**O hydrogen bonds. ^65^ However, it is increasingly recognized that C-H**…**O hydrogen bonds play an important role in molecular recognition processes^66^, protein folding stabilization^67^, in the interaction of nucleic acids with proteins^68^, in enzyme catalysis^69^, and in the stabilization of protein-ligand biding complexes.^70,71^ A matched pair of CDK2 inhibitors illustrates the contribution of C-H**…**O hydrogen bonds to protein-ligand complexes (Figure 7).^72^ The only difference between the two inhibitors is the substitution of a NH_2_ with a methyl group on the thiazole ring of compound **8** (Figure 7a). Although the N-H**…**O hydrogen bond of **7** is stronger than the C-H**…**O hydrogen bond of **8,** the latter compound is more potent probably due to the penalty associated with desolvating the NH_2_ of **7** upon binding.

**Figure 7.**
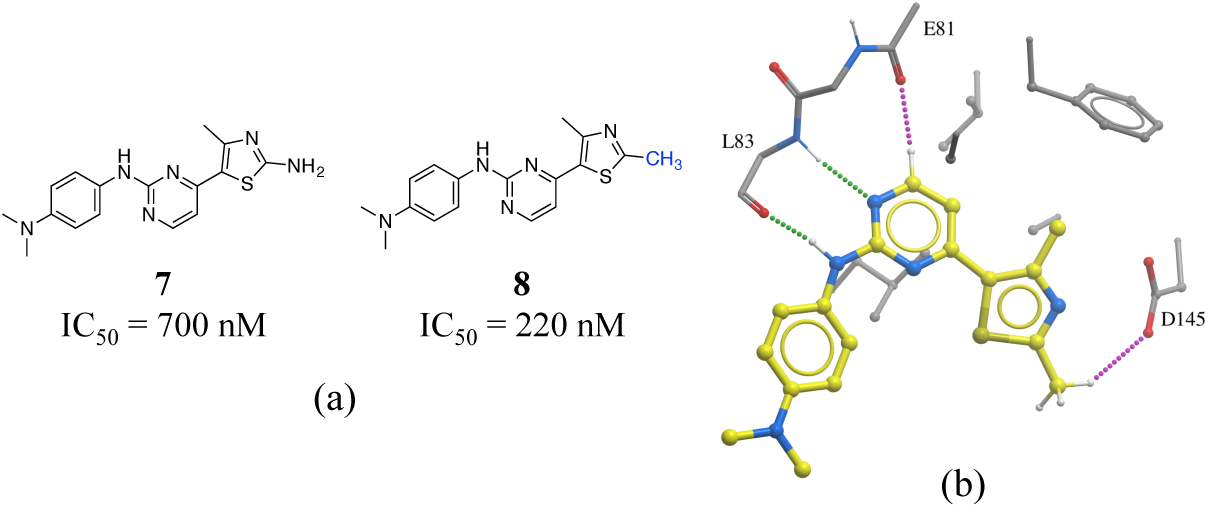
a) Chemical structure of two CDK2 inhibitors; b) X-ray cocrystal structure of human CDK2 and 8 (PDB: 1PXP). The N-H**…**O and C-H**…**O hydrogen bonds are displayed as green and magenta dotted lines, respectively.

### Salt bridges

The contact between a positively charged nitrogen and a negatively charged oxygen (i.e. *salt bridge*) was the fifth most frequent interaction type in our analysis (7276 interactions) (Figure 1). The number of salt bridge interactions with a positive nitrogen coming from the protein and the negative oxygen coming from the ligand was two times higher than the opposite (4882 *vs*. 2394 interactions, Table S2). This probably reflects the higher number of ligands containing carboxylic acids (1849) than ammonium groups (1103) in our dataset, as the frequency of arginine (5.6%) and lysine (5.0%) in proteins is similar to that observed for aspartic acid (5.4%) and glutamic acid (3.8%) (UniProtKB/TrEMBL UniProt release 2017_03).^73^ Arginine was the cation in 83.6% of all interactions (Figure S8). This seems to be agreement with quantum mechanical calculations, which predict that arginine are more inclined than lysine side-chains to form salt bridges.^74^ Finally, the distribution of negatively charged oxygens around the guanidinium group of arginine shows a higher density around the terminal (ω) nitrogens than at the secondary amine (ε) nitrogen (Figure S8).

Salt bridges contribute little to protein stability as the favorable binding energy obtained from forming a salt bridge is not sufficient to offset the energetic penalty of desolvating charged groups.^75,76^ However, the strength of salt bridge interactions is strongly dependent on the environment. In particular, buried salt-bridges can make crucial contributions to ligand binding.^77–79^ For example, the terminal N,N-dimethylamino tail of **10** forms a salt bridge with D831 in the kinase domain of epidermal growth factor receptor (EGFR) (Figure 8). When the nitrogen atom of the terminal N,N-dimethylamino group was replaced by a carbon (**9**) potency was reduced by more than 800-fold.^80^

**Figure 8.**
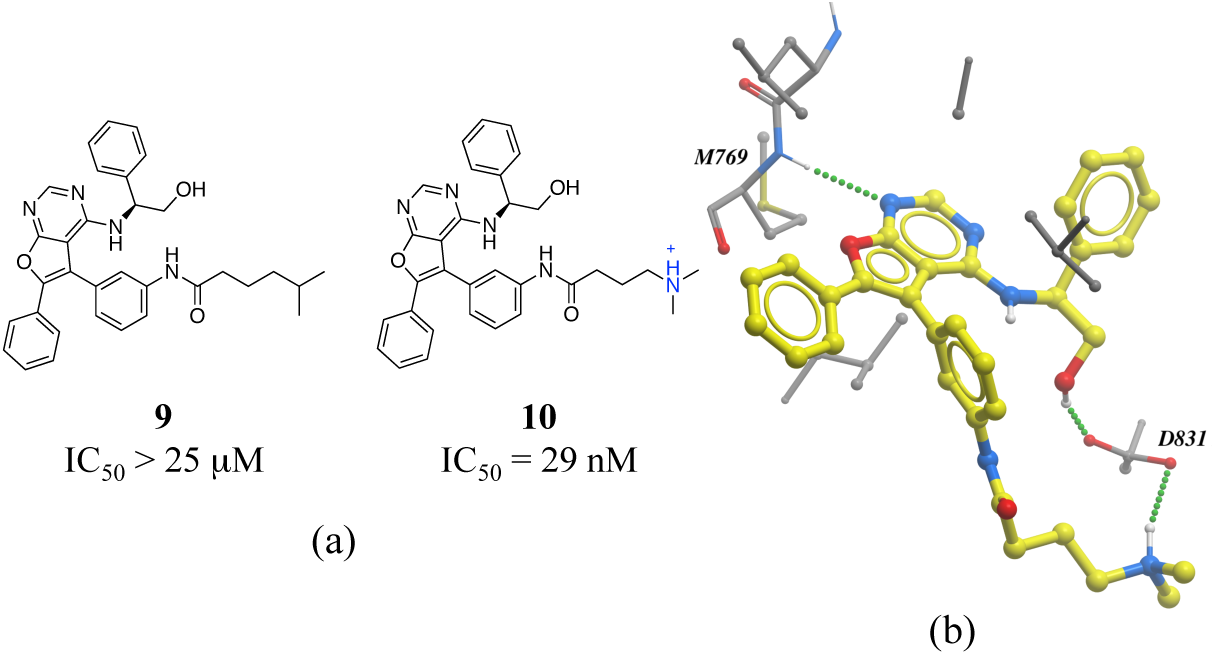
a) Chemical structure of two inhibitors of human EGFR; b) X-ray cocrystal structure of kinase domain of EGFR and **10** (PDB: 4JRV), the terminal N,N-dimethylamino tail of **10** forms a salt bridge with D831. Hydrogen bonds are displayed in dotted green lines.

### Amide…π stacking

Interactions between an amide group and an aromatic ring were the sixth most frequently observed (Figure 1). In these interactions, which are related to canonical aromatic π-stacking, the π-surface of the amide bond stacks against the π-surface of the aromatic ring.^81,82^ As previously observed for π-π stacking interactions, we did not find significant preference for face-to-face over edge-to-face arrangement (2907 and 2060 interactions respectively) (Table S2). The most frequent amino acids participating in face-to-face amide**…**π stacking were glycine (19.4%) and tryptophan (17.9%), while glycine (20.1%) and leucine (13.0%) were the most often observed in edge-to-face geometry (Figure S9). The fact that 88.5% of all amide**…**π stacking interactions occurred between the backbone amide group of a protein (generally a glycine) and the aromatic ring of a ligand points at a strategy to exploit peptide bonds in binding sites that is probably underused in structure-based drug design.

Amide**…**π stacking interactions are common and significant in protein structures.^83^ These interactions were also shown to sometimes play an important role in ligand binding.^84–86^ For example, the selective permeation of urea is facilitated by amide**…**π stacking interactions in the urea transporter.^87^ In another example, the 11-fold difference in *K*_*i*_ between a matched pair of oxazole-containing factor Xa inhibitors was attributed to the influence of the dipole of the oxazole ring on the amide**…**π interaction (Figure 9).^88^

**Figure 9.**
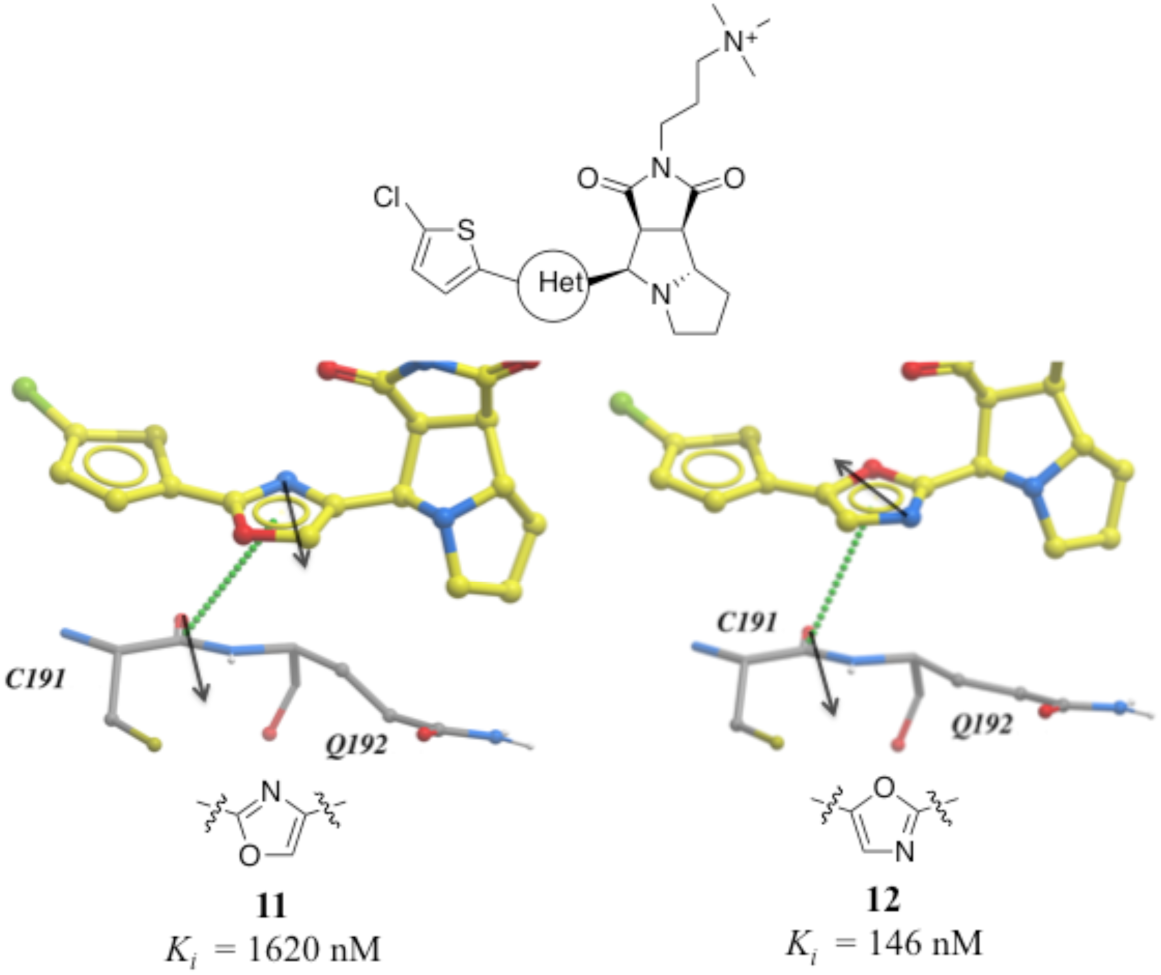
X-ray cocrystal structure of **11** (PDB: 2Y5H) and **12** (PDB: 2Y5G) bound at the active site of factor Xa. The amide**…**π stacking interaction is shown as dotted green lines. The dipoles of the oxazole ring and peptide amide (black arrows) are parallel in **11** and anti-parallel in **12**.

### Cation-π

We found 2577 interactions between a positively charged nitrogen and an aromatic ring (Figure 1). These cation-π interactions are essentially electrostatic due to the negatively charged electron cloud of π systems.^89^ In more than 90% of these interactions, the nitrogen came from the receptor and the aromatic ring from the ligand, reflecting, as previously noted, that drug-like compounds have often aromatic rings while ammonium groups are more rare (Table S2). Arginines were 3 times more frequently engaged in cation-π interactions than lysine side-chains. (Figure S10). A similar trend was previously observed for peptidic interactions.^90^ This preference has been attributed to the fact that the guanidinium group of arginines can donate several hydrogen bonds while simultaneously binding to an aromatic ring.^82^ When the positive nitrogen came from the ligand, tyrosine side-chains were the most common partner with 156 interactions, followed by phenylalanine and tryptophan (59 and 24 interactions respectively) (Figure S10). Potentiation of the cation-π binding ability of the tyrosine upon hydrogen bonding of its hydroxyl group was proposed to be at the origin of a similar bias in peptidic interactions.^90^

Cation–π interactions are widespread in proteins and are important determinants of the structure, stability, and function of proteins.^91^ An example that is especially compelling is the Royal family of epigenetic reader proteins, that feature an aromatic cage composed of two to four aromatic residues that make cation–π and hydrophobic interactions with postranslationally methylated lysines or arginines side-chains.^92^

Many drug-receptor interactions involve cation-π interactions. One of the earliest examples is the recognition of acetylcholine (ACh) by the nicotinic acetylcholine receptor (nAChR). Similarly, GABA^93^, glycine^94^, and 5-HT_3_^95^ receptors have all been shown to participate in cation–π interactions with neurotransmitters. In a series of insightful experiments, sequential methylation of an ammonium group in a series of potent factor Xa inhibitors gradually increased the binding affinity by 3 orders of magnitude.^96^ Comparing the affinity of a ter-butyl analog (compound **15**) with the trimethylated ammonium group (compound **17**), indicated that the cation–π interaction contributed to a 60 fold increase in potency (Figure 10).

**Figure 10.**
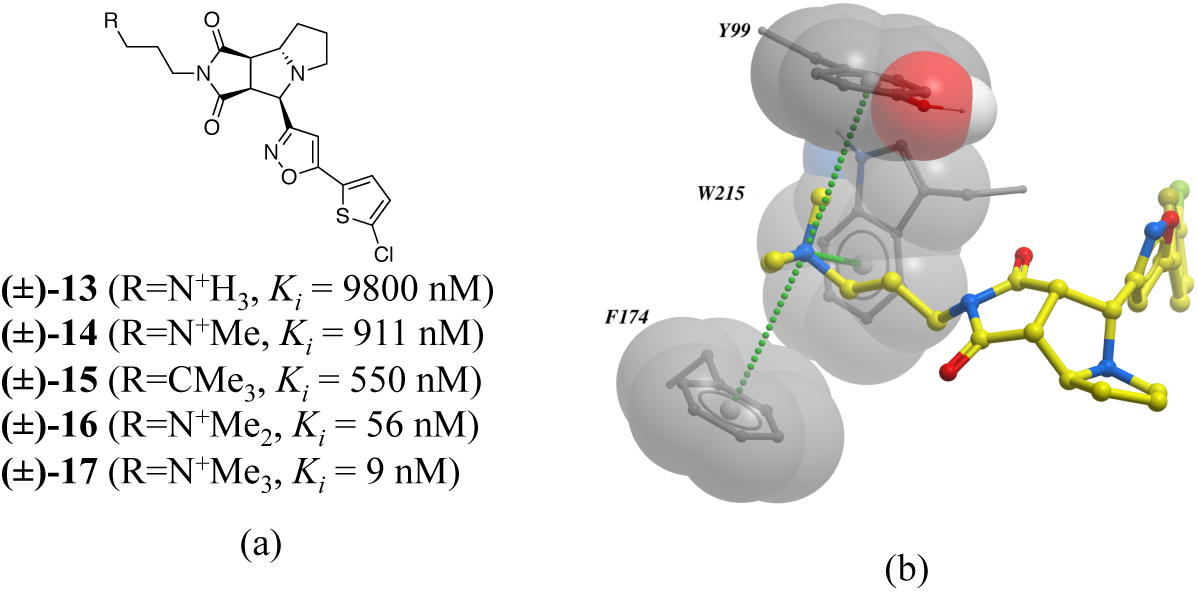
a) Chemical structure of a series inhibitors of human factor Xa; b) in the X-ray cocrystal structure of human factor Xa and **17** (PDB: 2JKH), the quaternary ammonium ion fill the aromatic box. The cation-π interaction is displayed as dotted green lines.

## Conclusion

We presented here a statistical analysis of the nature, geometry and frequency of atomic interactions between small molecule ligands and their receptors available in the PDB. The enrichment of polar interactions in bound fragments, but hydrophobic contacts in optimized compounds reflects the challenge of overcoming desolvation penalty during lead optimization. This unbiased census recapitulates well-known rules driving ligand design, but also uncovers some interaction types that are often overlooked in medicinal chemistry. This analysis will help in the interpretation of difficult SAR, and may serve as a knowledgebase for the improvement of scoring functions used in virtual screening.

## Acknowledgments

We thank Vijayaratnam Santhakumar for his helpful comments on this manuscript. The SGC is a registered charity (number 1097737) that receives funds from AbbVie, Bayer Pharma AG, Boehringer Ingelheim, Canada Foundation for Innovation, Eshelman Institute for Innovation, Genome Canada through Ontario Genomics Institute [OGI-055], Innovative Medicines Initiative (EU/EFPIA) [ULTRA-DD grant no. 115766], Janssen, Merck & Co., Novartis Pharma AG, Ontario Ministry of Research, Innovation and Science (MRIS), Pfizer, São Paulo Research Foundation-FAPESP, Takeda, and the Wellcome Trust.

## Table of Contents Graphic

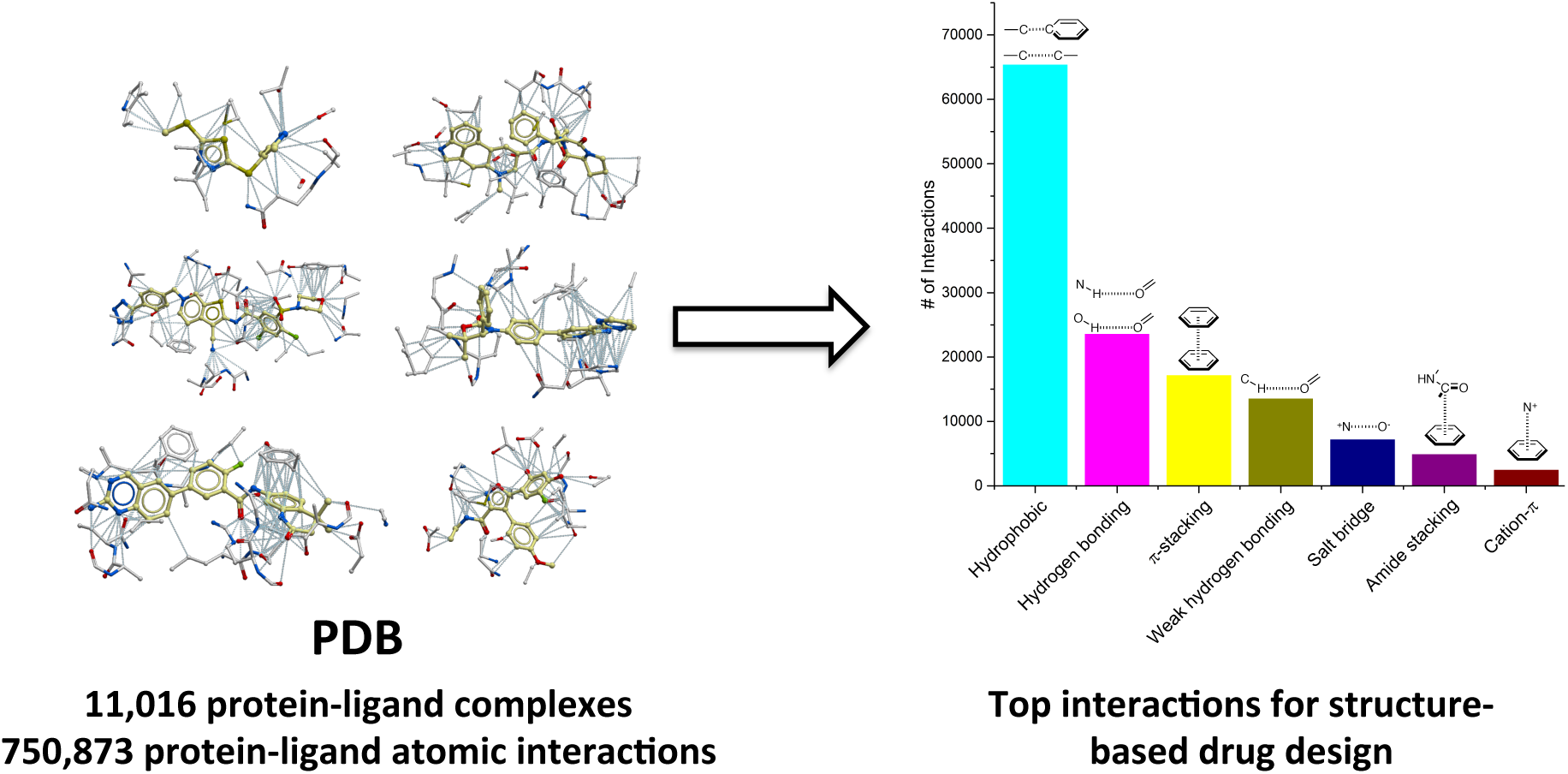

